# Bacterial cell widening alters periplasmic size and activates envelope stress responses

**DOI:** 10.1101/2022.07.26.501644

**Authors:** Matylda Zietek, Amanda Miguel, Iskander Khusainov, Handuo Shi, Abir T Asmar, Sri Ram, Morgane Wartel, Anna Sueki, Martin Schorb, Mark Goulian, Jean-François Collet, Martin Beck, Kerwyn Casey Huang, Athanasios Typas

## Abstract

The Rcs signal transduction system is a phosphorelay responsible for sensing a wide variety of enterobacterial cell envelope stresses. In *Escherichia coli*, the Rcs system is required to survive A22 and mecillinam treatment, two drugs that perturb cell size. To test whether cell size changes might be correlated with envelope damage and thereby sensed by the Rcs system, we tuned *E. coli* cell size via drug inhibition with A22, point mutations to the cell-shape determinant MreB, and mechanically confined growth. In all conditions, cell width was strongly correlated with Rcs activation, with wider cells exhibiting more activation than wild-type. In all conditions, RcsF, the outer membrane-localized upstream component of the Rcs system, was essential for responding to cell width changes. Consistently, several envelope gene deletions known to induce the Rcs system via RcsF resulted in cells that were wider than wild-type. Cryo-electron microscopy revealed that the periplasm of a wide MreB mutant was on average ∼3 nm thinner than wild-type, thereby bringing RcsF closer to the downstream components of the signaling cascade in the inner membrane. Conversely, extending the flexible linker region of RcsF by ∼3 nm increased Rcs activity in wild-type cells. In summary, we propose that the Rcs system responds to changes in cell width because of altered periplasmic thickness.

## Introduction

Although bacteria have the capacity to respond to environmental changes and stresses through signal transduction pathways, in many cases how such changes or stresses are directly sensed remains unclear. In the case of defects in cell envelope integrity or assembly in Gram-negative bacteria, damage is often sensed at the cell surface [1]. Signals are then transduced to the cytoplasm via phosphorylation to activate the expression of repair and adaptive systems, and thereby maintain viability [2]. There are several signaling systems that have been reported to respond to stress in the Gram-negative cell envelope [3, 4].

One of the most well-studied and complex signaling pathways involved in responding to envelope stress is the Rcs phosphorelay, which responds to both outer membrane (OM)- and peptidoglycan-related stresses in enterobacteria [5]. Unlike typical two-component systems consisting of only a sensor histidine kinase and a cytoplasmic response regulator, the Rcs system has at least 4 additional components, including an intermediate inner membrane (IM) phosphorelay protein, an auxiliary non-phosphorylatable transcription factor, and two proteins that act upstream of the phosphorelay cascade and are associated with signal sensing (IgaA and RcsF). The OM lipoprotein RcsF, the most upstream component of the pathway, is required to activate the Rcs phosphorelay under most conditions. RcsF senses envelope damage by monitoring activity of the Bam machinery [6]. In growing cells, BamA, the major component of the outer membrane porin (OMP) assembly machinery, continuously funnels RcsF through OMPs to the cell surface [6, 7]. This process spatially separates RcsF from IgaA, which inhibits the Rcs pathway [8, 9]. The Rcs system is activated when BamA fails to bind RcsF and funnel it to OMPs, whereupon newly synthesized RcsF remains facing the periplasm and directly interacts with IgaA to activate the cascade [6, 10]. RcsF can also directly sense lipopolysaccharide (LPS) defects on the OM surface via a set of charged residues [6, 11–13]; activation presumably occurs via RcsF release and/or immobilization facing the periplasm. Interestingly, RcsF signaling is sensitive to periplasmic thickness (i.e. the distance between the OM and IM). When peptidoglycan becomes untethered from the OM, RcsF cannot reach IgaA to activate the system [1]. How RcsF orientation, periplasmic thickness, and other physiological parameters such as cellular dimensions are coupled to envelope damage has yet to be determined.

A chemical genomics screen of the *E. coli* Keio library of nonessential gene knockouts [14] revealed that cells lacking components of the Rcs pathway (Δ*rcsB*, Δ*rcsD*, and Δ*rcsF*) display increased sensitivity to specific conditions that perturb the cell envelope [15]. These conditions included the MreB inhibitor A22 [16], the PBP2 inhibitor mecillinam [17], the osmolyte NaCl, and the LPS biosynthesis inhibitor CHIR-090 [18, 19]. Under each of these conditions, the Rcs pathway becomes active in wild-type cells in an RcsF-dependent manner [6]. During A22 and mecillinam treatment, cells also become rounder by increasing in width and decreasing in length [20], but whether changes in cellular dimensions can lead to activation of the Rcs pathway remains unknown.

Bacterial cells possess mechanisms to directly sense local biophysical properties and architectural changes such as surface curvature [21], envelope stress [22], and periplasmic thickness [1]. However, there is currently no bacterial signaling system known to intrinsically sense changes in cellular dimensions and/or shape. Cell shape, which varies across bacterial species [23], is typically tightly maintained by a given species in a non-fluctuating environment [24]. Yet changes to cell shape can have important physiological consequences, such as during infection in which smaller and rod-shaped cells are better at escaping the host immune response than larger and rounder cells [25]. Importantly, cell shape and dimensions are largely controlled by cell wall and OM biosynthesis [26, 27]. Hence, we wondered whether the Rcs system directly senses changes in global cellular dimensions.

Here, we report a general relationship between the Rcs stress response and cell width. We find that the Rcs system is activated under environmental, genetic, and mechanical perturbations to cell width. Cryo-electron microscopy of wider *E. coli* mutant cells revealed significant decreases in periplasmic thickness relative to wild-type cells, providing an explanation for increased Rcs activation by shifting RcsF closer to its IM interaction partner, IgaA. Consistent with this hypothesis, in cells with wild-type periplasmic thickness, extending the RcsF linker resulted in Rcs activation. These findings support a model in which Gram- negative bacteria use RcsF to sense and respond to cell-envelope rearrangements associated with cell widening.

## Results

Rcs activation by cell shape-perturbing antibiotics is correlated with cell width To interrogate Rcs activation and its coupling to cellular dimensions, we used *E. coli* DH300 cells, an MG1655 Δ*argF-lac* strain with *lacZ* reintegrated on the chromosome under control of the promoter of *rprA*, a small RNA expressed only during Rcs activation [28]. We monitored β-galactosidase production and cellular dimensions upon treatment with 5 µg/mL A22. Approximately 15 min after the start of A22 treatment, β-galactosidase activity started to increase in wild-type DH300, but not in Δ*rcsF* cells (Fig. 1A); note that synthesis and folding of β- galactosidase requires only a few minutes [29]. The delayed Rcs activation coincided with the timing of changes in mean cell width, while cell length started to gradually decrease immediately after A22 treatment (Fig. 1B). Rcs activity and mean cell width increased in an A22 dose-dependent manner (Fig. S1A). As expected, A22-induced changes in mean cell width were not Rcs-dependent (Fig. 1B). Moreover, A22 treatment did not affect growth rate, at least for the first 60 min (Fig. 1A, S1A), as previously reported [30].

**Figure 1:**
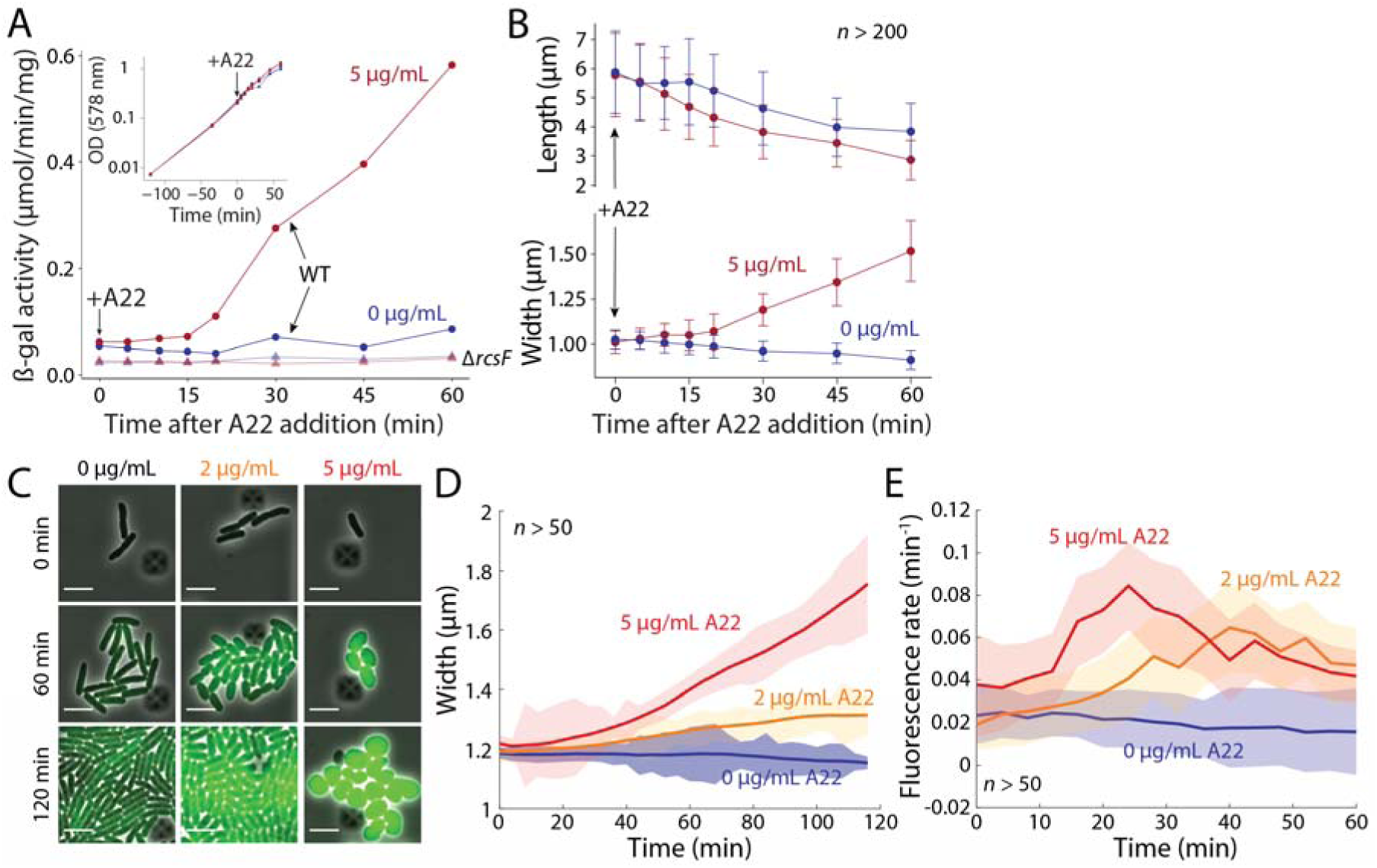
A22 concentration-dependent activation of the Rcs pathway is correlated with cell width changes. A) The Rcs system was activated ∼15 min after addition of 5 µg/mL A22, as measured by following the induction of chromosomal *rprA::lacZ* in a β- galactosidase assay. Cells were treated with A22 at OD_578_=0.3. Inset: growth was unaffected by A22 addition. The Rcs system was not activated in Δ*rcsF* cells (triangles). Data are a representative of four biological replicates (see Fig. S1B for other replicates). B) Cell length gradually decreased, starting immediately after A22 addition, while width started to increase ∼15 min after A22 addition, a similar delay as for Rcs activation in (A). Curves are means and error bars represent 1 standard deviation (*n*>200 cells for every time point). Cells were sampled and fixed from the same experiment shown in (A). C) Time-lapse images of typical *E. coli* cells expressing msfGFP from the Rcs- dependent *rprA* promoter on plasmid pMZ13 (Methods) treated with various concentrations of A22. D) Mean cell width of the populations represented in (C) increased over time, faster for higher A22 concentration. Curves are means and shaded regions represent 1 standard error of the mean (SEM, *n*>50 cells for every time point); the SEM was occasionally smaller than the width of the lines. E) Rcs activation increased in an A22 dose-dependent manner. Curves are means of rate of msfGFP fluorescence production and error bars represent 1 SEM (*n*>50 cells for every time point); the SEM was occasionally than the width of the lines. Even untreated cells exhibited a basal slow rate of msfGFP production.

To concomitantly monitor Rcs activation and morphology in single cells, we transformed wild-type cells with a low-copy plasmid that expresses msfGFP from the *rprA* promoter (Methods) and performed time-lapse imaging during A22 treatment (Fig. 1C). As expected, *rprA* expression was RcsF-dependent. Cell width increased more (Fig. S2A) and at a faster rate for higher A22 concentrations (Fig. 1D). As in our population-based data, upon treatment with 5 µg/mL A22, a short (10-20 min) delay was evident before the rapid width increase and Rcs activation (Fig. 1D,E). Normalized msfGFP levels (total intensity divided by cell volume) initially increased at an A22 dose-dependent rate in cells (Fig. 1E), although variation in cell width within a population of cells did not consistently correlate with *rprA* expression (Fig. S2B). Taken together, these data suggest that cell width increases may be sufficient for Rcs activation.

### Mutations to MreB cause correlated changes to cell width and Rcs activation

To test whether genetically induced cell-width increases would activate the Rcs pathway similarly to A22 treatment, we transformed *E. coli* MG1655 MreB point mutants that exhibit a range of mean cell widths (Fig. 2A) [31] with the *rprA* msfGFP-reporter plasmid; these mutants have the same maximal growth rate as cells with the wild-type *mreB* allele [31]. We imaged each of these strains during steady-state exponential growth, and quantified msfGFP intensity using single-cell microscopy. Average msfGFP fluorescence was strongly correlated with cell width and deletion of *rcsF* eliminated fluorescence in all strains (Fig. 2B), suggesting that increased cell width generally activates the Rcs pathway in a RscF-dependent manner.

**Figure 2:**
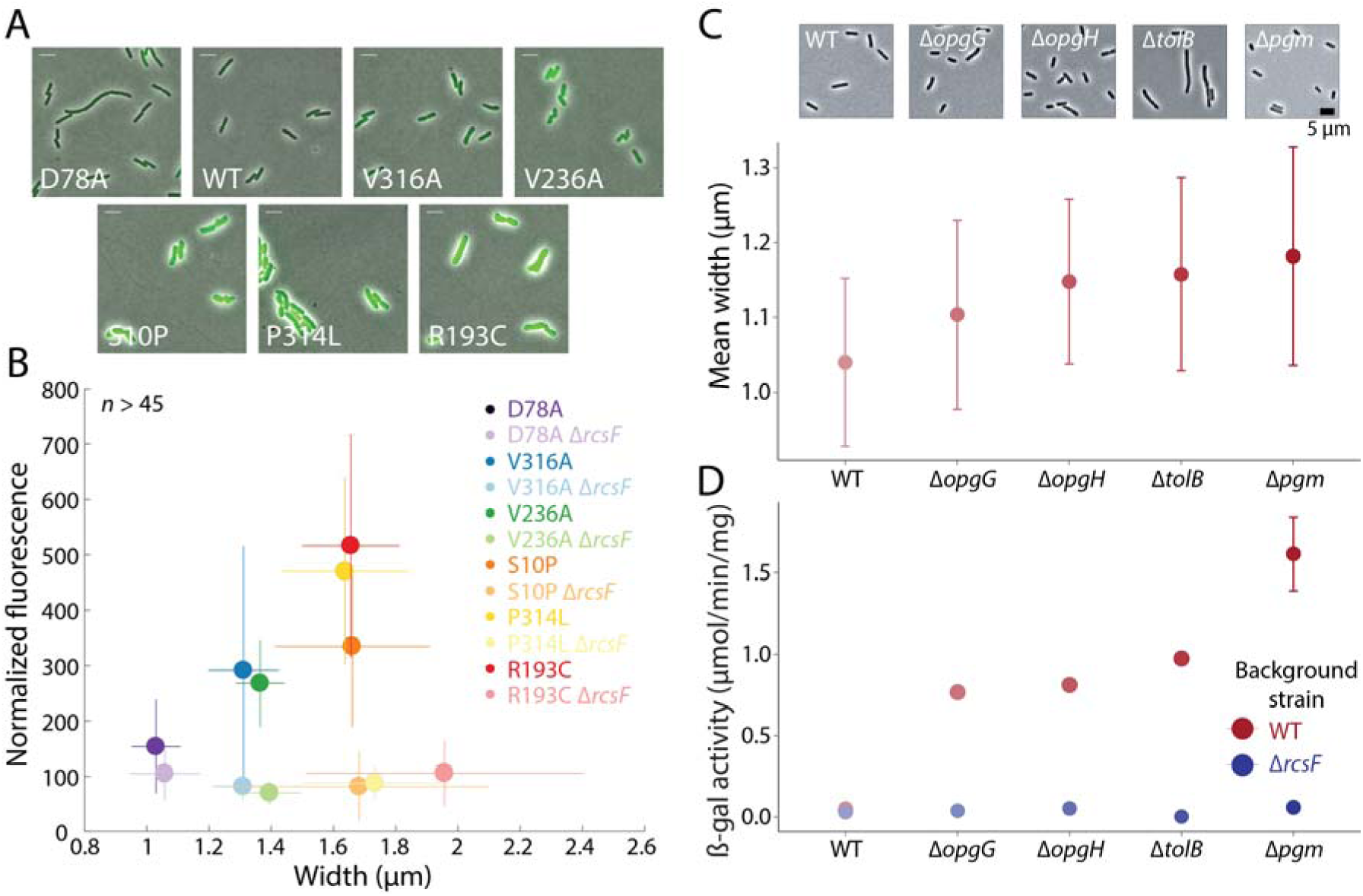
Rcs activation is correlated with cell width in mutants with altered cell size. A) Overlay of phase-contrast and msfGFP fluorescence from the Rcs- dependent *rprA* promoter in MreB mutants with a range of cell widths. B) msfGFP intensity normalized to cell volume was highly correlated with cell width across the MreB mutants in (A). Fluorescence remained low in all mutants when *rcsF* was deleted. Data points are mean values and error bars represent 1 standard deviation. *n*>45 cells for each strain. C) Representative images (top) and mean cell width (bottom) of cells from various single-gene deletion mutants known to activate the Rcs system. Cells were harvested during exponential growth in LB and fixed. Circles are means and error bars are 1 standard deviation (*n*>1100 cells per strain). D) Rcs pathway activity as measured by β-galactosidase of the *rprA::lacZ* fusion in the same mutants and conditions as in (C). Rcs activity was higher in wider deletion mutants. Circles are means and error bars are 1 standard deviation (*n≥*3 replicates per strain). The error bars for all strains besides Δ*pgm* are smaller than the circles.

### Genetic perturbations that induce the Rcs pathway also affect cell width

To determine how closely cell width is connected with Rcs activation, we examined the cell dimensions of envelope mutants that are known to activate the Rcs system (Δ*opgG*, Δ*opgH*, Δ*tolB*, Δ*pgm* [32]) (Fig. 2C). These mutants perturb the cell envelope in different ways, with *opgG* and *opgH* encoding osmo-regulated periplasmic glucans [33], *tolB* encoding a key component of the Tol-Pal trans-envelope machinery that regulates OM constriction and homeostasis [34–36], and *pgm* encoding a phosphoglucomutase that has been implicated in cell division [37, 38]. Each of these mutants had a larger mean cell width than wild type (Fig. 2C). Moreover, Rcs activation was dependent on RcsF and scaled with width changes (Fig. 2D), supporting a general connection between width and Rcs activation.

### Mechanical deformation leads to Rcs activation

In a previous study, *E. coli* cells placed under mechanical confinement through a compressing membrane gradually grew into pancake shapes and increased in cell width (as measured in the imaging plane) [39]. To test whether such mechanically induced width changes would activate RcsF similarly to chemical and genetic perturbations, we subjected wild-type and Δ*rcsF* cells carrying the *rprA* GFP-reporter to (non-uniform) mechanical confinement (Fig. 3A). Cells grew into a wide range of morphologies due to local changes in the degree of confinement in our device (Fig. 3B). Interestingly, Rcs activation was RcsF- dependent and correlated with the minor axis of cells (width) (Fig. 3C). Thus, mechanically induced width changes also result in RcsF-dependent Rcs activation.

**Figure 3:**
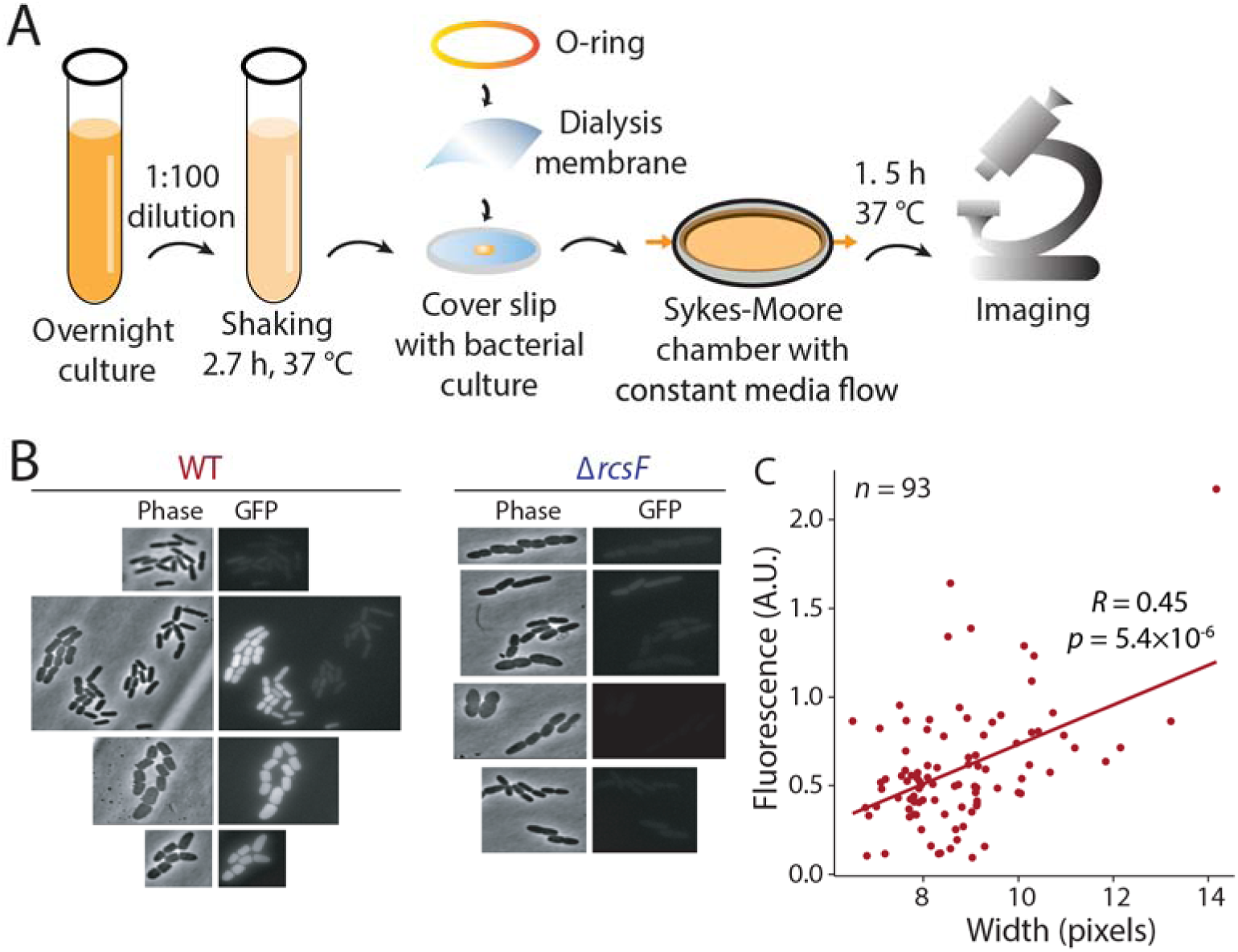
Mechanical perturbations activate the Rcs pathway. A) Schematic of protocol for monitoring cell growth under mechanical confinement. Log-phase cells carrying an *rprA*::GFP fusion on plasmid pP_rprA-gfp and constitutively expressing mCherry from the chromosome were placed under a permeable membrane and pulling forces were exerted on the membrane edge with a Sykes-Moore chamber to reduce the volume under the membrane. In these conditions, cells grew in different shapes depending on the differential confinement forces exerted across the membrane surface. Cells were imaged after 1.5 h. B) Representative images of an experiment as described in (A). Upon mechanical confinement, wild-type and Δ*rcsF* cells increased similarly in major- and minor-axis lengths (proxies for cell length and width, respectively). On the other hand, GFP intensity increased only in wider wild-type (WT) cells, but not in any Δ*rcsF* cells. In some fields of view, wild-type cells with both increased and normal cell width were visible; only the wider cells exhibited msfGFP expression. C) The ratio of GFP to mCherry intensity was correlated with cell width in wild-type cells.

### Wider MreB mutant exhibits decreases in periplasmic thickness sufficient to activate the Rcs pathway

To probe the mechanism of Rcs activation in wider MreB mutants, we first tested whether RcsF transport from the IM to the OM was disrupted. RcsF remaining in the IM leads to constitutive activation of the Rcs system [6, 40]. However, RcsF localization to the OM was intact in the widest mutant MreB^R193C^ (Fig. S3). A previous study showed that transmission of damage signals from the cell envelope by RcsF, located in OM, to the IM-localized IgaA depends on the distance between the two membranes relative to the length of the RcsF flexible linker [1]. We thus wondered whether cell widening may alter periplasmic thickness in a manner that allows RcsF to reach IgaA more readily and activate the Rcs system.

Since the Rcs pathway was highly active in wide MreB mutants during normal growth (Fig. 2B), we sought to determine whether MreB^R193C^ cells had different periplasmic thickness from wild-type cells. Using cryo-electron microscopy, we acquired high-resolution images of wild-type and MreB^R193C^ cells. We focused on the midsection of cells, which exhibited more uniform envelope architecture (Fig. 4Ai), computationally segmented the IM and OM (Fig. 4Aii,iii; Methods), and measured the intermembrane distance at regularly spaced points as a proxy for periplasmic thickness. (Fig. 4Aiv, S4). The wider MreB mutant exhibited a significant decrease by ∼3 nm in periplasmic thickness (Fig. 4B, S5). This decrease in periplasmic thickness should mean that newly synthesized RcsF reaching to the OM should be closer to the IM-located IgaA, which would facilitate their interaction. Consistent with this idea, extending the RcsF linker by 7 amino acids (2-3 nm, RcsF^+7^) in wild-type cells resulted in Rcs activation (Fig. 5A). The RcsF^+7^ construct retained its outer membrane localization (Fig. 5B), demonstrating that it was indeed the increase of the RcsF length relative to the intermembrane distance in MreB^R193C^ cells that led to Rcs activation. Thus, changes in periplasmic thickness due to cell widening trigger the Rcs system.

**Figure 4:**
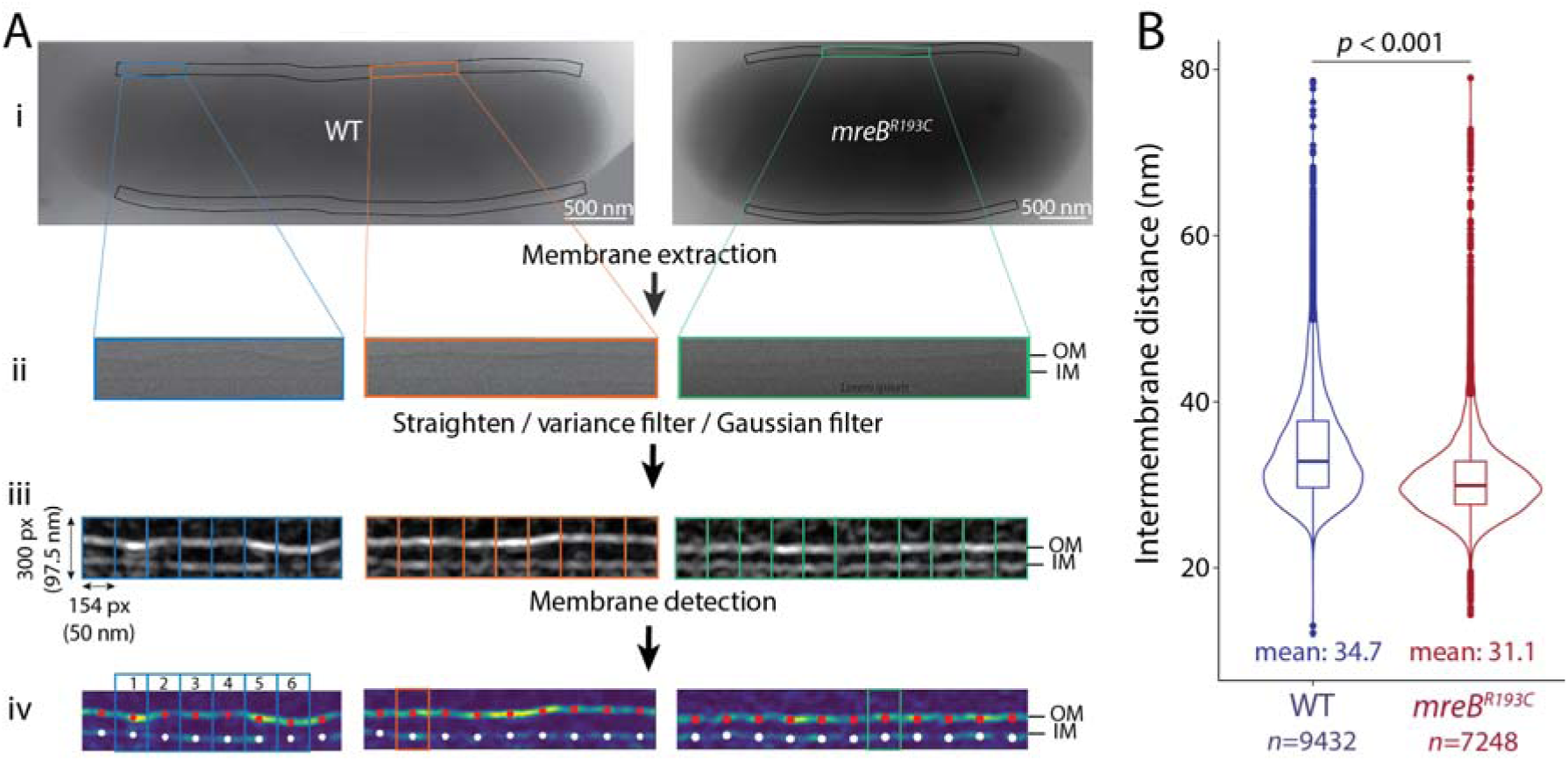
Periplasmic thickness is decreased in wider *mreB^R193C^* cells compared with wild-type. A) Blended montage of a stack of cryo-EM images of representative cell with average intermembrane distance for wild-type (left) and mreB^R193C^ (right). Measurements were carried out along cylindrical regions excluding the tips (black boxes in (i)). Both wild-type and *mreB^R193C^* cells exhibited diverse membrane morphologies from straight (orange, green) to wrinkled (cyan). The processing workflow involved membrane extraction (ii), membrane enhancement via straightening and filtering (ii), and separation into 50-nm segments (iii). Measurements were computed as the distance between the red and white dots in each segment (Fig. S4). OM, outer membrane; IM, inner membrane. B) Intermembrane distances were lower on average in *mreB^R193C^* cells compared with wild-type. Violin plot shows the smoothened density of measurements, and the boxplot shows the first quartile, median, and third quartile. *n* represents the number of points at which measurements were made, for 86 wild-type and 71 *mreB^R193C^* cells from three biological replicates.

**Figure 5:**
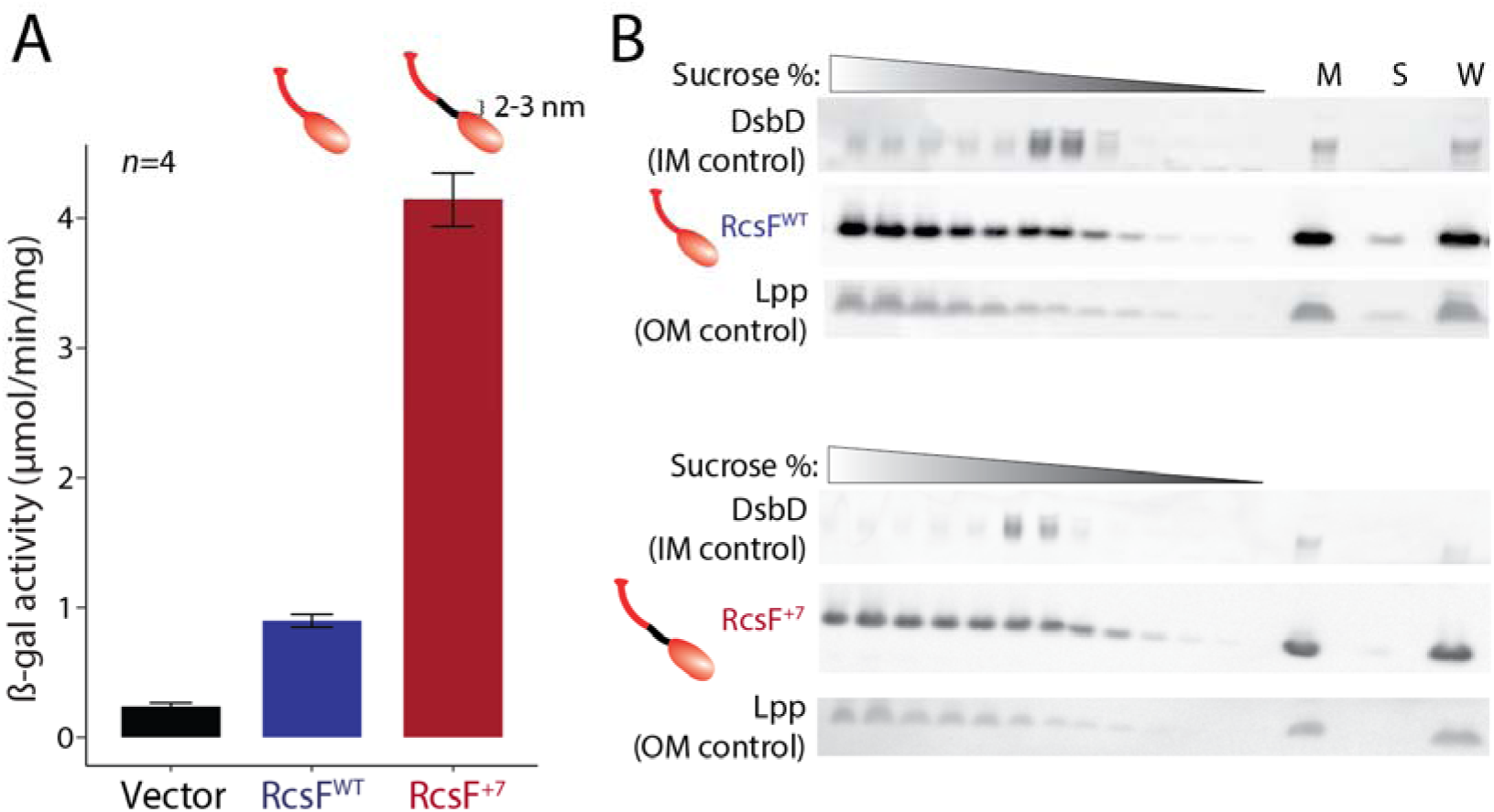
Extension of the RcsF linker results in Rcs activation in wild-type cells without affecting RcsF localization. A) Activation of the Rcs pathway as measured by β-galactosidase activity was higher in a mutant expressing RcsF with its linker region extended by 7 amino acids (RcsF^+7^) cells compared to wild-type or the vector control in Δ*rcsF* cells. Bacteria were harvested for β-galactosidase assay at OD_600_=0.6, error bars represent 1 standard deviation. B) Sucrose gradient fractionation of cells expressing RcsF^WT^ or RcsF^+7^ showed similar localization of both proteins to the outer membrane (OM). Immunoblots of RcsF and controls (DsbD for the inner membrane (IM) and Lpp for the OM). M, total membrane sample prior to fractionation; S, soluble non-membrane fraction; W, whole lysate.

## Discussion

Our data support a strong correlation between width and Rcs activation, but does cell width directly activate the Rcs pathway? In the case of mechanical squashing, the width variation across the population due to heterogeneity in the application of force was correlated with *rprA* expression (Fig. 3C). *rprA* expression also scaled with mean cell width at the population level across A22 concentrations (Fig. 1D,E) or cell-width mutants (Fig. 2B,D), although within-population variability in width at fixed A22 concentration was not consistently correlated with *rprA* expression (Fig. S2B), presumably due to the relative minor fluctuations of cell width and fluorescence signal reporting on Rcs activation with some delay. However, despite the many conditions reported here in which cell width and Rcs activation were strongly correlated, the Rcs system can also be activated in conditions in which cell width is unaltered. We previously reported that malfunctions in the Bam machine (e.g., lower BamA levels) lead to Rcs activation [6], because RcsF cannot be funneled to the cell surface and remains facing the periplasm. Similarly, defects in LPS can transiently activate the Rcs system [11, 12], and Δ*rcsF* cells are sensitive to CHIR-090 treatment [15] even though CHIR-090 activates the Rcs pathway in wild-type cells but does not change cell width (Fig. S6A). Deletion of *rfaF*, which encodes an enzyme involved in LPS core biosynthesis [41], resulted in Rcs activation without affecting cell width (Fig. S6B).

Taken together, width changes are sufficient but not necessary for Rcs activation, and width is correlated with another feature(s) that ultimately dictates Rcs activation. Indeed, our data implicates periplasmic thickness as a key factor in Rcs activation, which was previously shown to be affected by the presence and/or length of the lipoprotein Lpp and to dictate communication across the envelope by RcsF to IgaA [1]. Extension of the RcsF linker has been shown to compensate for increases in periplasm thickness in *lpp* mutants [1]. Here, extension of the RcsF linker by 2-3 nm was sufficient to activate the Rcs system in wild-type cells (Fig. 5A), to a similar extent (4- to 5-fold) as Rcs was activated in MreB^R193C^ cells (Fig. 2B). Extension of the linker or decrease of the periplasmic thickness brings RcsF closer to the low abundant, integral IM protein, IgaA (∼220 copies/cell) [42], with which RcsF binds with high affinity (*K*_D_∼1.6 nM) [43]. Consistent with the hypothesis that the Rcs system is activated by a mismatch between RcsF size and periplasmic thickness, our cryo-electron microscopy data indicated that MreB^R193C^ cells had a ∼3 nm thinner periplasm compared with wild-type cells (Fig. 3B).

How cell shape and size are connected to periplasmic dimensions remains mysterious. One possibility is that cell widening is correlated with changes in turgor that cause the periplasm to become thinner. Unfortunately, periplasmic size is essentially unknown in all but a few cases. Cryo-EM provides a means to quantify periplasmic dimensions in a native state, but our measurements highlight the high variability in periplasmic thickness both across cells and within cells along the cell surface (Fig. 4B, S5), which has previously been underappreciated [1]. This heterogeneity in periplasmic thickness suggests that there may be the potential for Rcs activation in every cell dependent on the localization of RcsF and its associated machinery, and motivates the development of methods that i) can quantify periplasmic dimensions at the poles, which are difficult to discern without tomography; and ii) directly measure Rcs activation at a subcellular level. Importantly, while our data suggests that cell width and periplasmic thickness are correlated, defects in the Bam machinery and LPS can also activate the Rcs system, presumably without affecting periplasmic size. Newly transported RcsF in the OM is ushered to the cell surface by the Bam machinery, and interactions with BamA (*K*_D_∼400 nM; 3,900 copies/cell) [7, 42] and with the highly abundant OmpA (*K*_D_∼100 µM; 100,000-200,000 copies/cell) [42–44] prevent RcsF from reaching to IgaA and activating the Rcs signaling system. Hence, akin to alterations in cell width, alterations to periplasmic thickness are sufficient but not necessary for Rcs activation.

Independent of the mechanism of activation, the role of Rcs activation upon changes to cellular dimensions is intriguing. Periplasmic glucans, which are induced by the Rcs pathway [32], may themselves physically determine periplasmic thickness, as deletion of the genes responsible for their production activates Rcs [45] and increases cell width (Fig. 2D). Rcs activation as cells widen may serve to protect from increased incidence of envelope stress (e.g. protein misfolding and mislocalization [8, 32]) and/or greater sensitivity to rupture due to changes in the distribution of mechanical stresses in the envelope, and may be involved in width-dependent feedback on *mreB* transcription [27]. In particular, Rcs-induced production of colanic acid [46] could create an additional layer of mechanical strength to protect the cells from rupture, similar to how the OM contributes to sustaining the internal turgor pressure [47]. Finally, given the recently discovered connections between cell width and length through surface area-to-volume ratio [48, 49], the width-dependence of Rcs activation suggests that the Rcs pathway may play an important role in cell division and length determination. Interestingly, the Rcs response has been reported to directly activate *ftsAZ* expression [50], is required for *de novo* envelope biogenesis [51], and leads to non-growing cells upon constitutive activation (*igaA* depletion [6, 52]). In an accompanying paper, we uncouple cell-shape sensing of the Rcs system from its response and characterize the effects of ectopic and constitutive Rcs activation on cell growth and shape. Together, these studies highlight the intimate connections between cellular structure and stress sensing.

## Methods

### Strains and plasmids

All bacterial strains and plasmids used in this study are listed in Table S1 and S2, respectively. The *E. coli* K12 strain MG1655 derivative, DH300 [28], was used as the main genetic background unless otherwise mentioned, and is referred to as wild-type. Rcs activity was monitored by either β-galactosidase assays using a chromosomal transcriptional fusion of the Rcs-dependent promoter of *rprA* to *lacZ* [28], or by fluorescence microscopy using a plasmid expressing P*rprA::sfGFP (*pMZ13). Gene deletion strains (Δ*rcsF::kan*, Δ*opgG::kan*, Δ*opgH::kan*, Δ*tolB::kan*, Δ*pgm::kan*) were created by P1 transduction of the corresponding Keio library mutant [14] into the DH300 background. All deletions were verified by PCR/Sanger sequencing. When necessary, antibiotic markers were flipped out [53]. MreB mutants were created in MG1655 as previously described [31]. Deletion of *rcsF* from the MreB mutants was accomplished via P1 transduction from the Δ*rcsF* Keio strain. pRcsF^WT^ and pRcsF^+7^ plasmids were described in [1].

The pMZ13 plasmid was constructed via Gibson assembly from pP_rprA-gfp, a derivative of pAC581 [54], which has copy number ∼15 and contains codon-optimized *gfpmut3* expressed from the *rprA* promoter. GFPmut3 was replaced with monomeric super-folder GFP (*E. coli* codon-optimized with the V206K mutation), which was PCR-amplified from pSFGFP-N1 (Addgene) [55].

The strain (MMR60) used for mechanical confinement contains the pP_rprA-gfp plasmid and *mcherry* expressed from the chromosome for normalization of the GFP signal [56].

### Growth conditions for various experiments

Unless otherwise stated, for single-cell time-lapse imaging, cells were grown in LB Lennox at 37 °C overnight, then diluted 1:200 and grown for 1.5 h. When appropriate, drugs were added to the culture and cells were incubated for an additional 30 min. Cells were then placed on agar pads and imaged for 60 min.

For probing *rprA*::*lacZ* activity in knockout strains (Fig. 2D), cells were grown overnight in LB at 37 °C, then diluted to an OD_578_ of 0.0075. Rcs system activity was measured at OD_578_=0.5 by β-galactosidase assay using a standard protocol [57]. For A22 experiments (Fig. 1A, S1), cells were grown in LB Lennox at 37 °C overnight, then diluted to an OD_578_ of 0.0075. A22 was added when OD_578_ reached 0.1 or 0.3 and Rcs system activity was monitored for 60 min by measuring β - galactosidase activity.

For comparing RcsF^WT^ and RcsF^+7^ activity (Fig. 5A), β-galactosidase assays were performed as described in [1].

### Single-cell imaging

For batch-culture experiments, cellular dimensions were examined in parallel with *rprA* promoter activity. For each time point, 1 mL of cell culture was harvested. Cells were fixed by incubation with 4% formaldehyde for 10 min at room temperature and centrifuged for 5 min at 1000*g* before being washed 3 times with 1 mL PBS. Next, cells were immobilized on 1% agarose pads and imaged with a Nikon Eclipse Ti inverted microscope, equipped with a Nikon DS- Qi2 camera and a Nikon Plan Apo Lambda 60X oil Ph3 DM phase-contrast objective. Images were acquired with NIS-Elements v. AR4.50.00 and analyzed using *Morphometrics* [58] and custom Matlab scripts.

For experiments examining cell width and *rprA::msfGFP* expression under A22 treatment (Fig. S2), cells were diluted 1:5000 from an overnight culture. At OD=0.4, cells were diluted 1:200 in 0, 0.25, 0.5, 1, 2, or 5 µg/ml A22 and incubated for 2.5 h before imaging on an agarose pad.

For time-lapse experiments under A22 treatment (Fig. 1C-E), cells were back-diluted 1:5000 from an overnight culture. At OD=0.2, cells were back diluted 1:10 onto LB pads made with 1% agarose and 0, 0.25, 0.5, 1, 2, or 5 µg/ml A22. Cells were imaged under phase contrast and fluorescence for 2 h.

### Image analysis

Images were analyzed using *Morphometrics* [58] and custom Matlab scripts. Time-lapse data was preprocessed using the machine learning segmentation software *DeepCell* [59] with manually curated training datasets specific to the microscope used for imaging. The contour outputs from *DeepCell* were then processed normally through *Morphometrics*.

### Microfluidics

For single-cell tracking in CellAsic microfluidic flow cells, cells were back-diluted 1:500 and grown for 3.5 h. Cells were then diluted to OD∼0.001 and placed in a CellAsic B04A chamber. Once introduced into the imaging chambers, cells were allowed to grow in LB for 20 min before switching to LB supplemented with A22 and were imaged for >4 h.

### Mechanical confinement using a Sykes-Moore chamber

An overnight culture of *rprA*-GFP cells grown in Minimal A medium with 0.2% glucose, 1 mM MgSO_4_, and 20 µg/mL chloramphenicol was diluted 100-fold and placed in a roller drum at 37 °C for 2.7 h. Five microliters of this culture (OD∼0.1) were pipetted onto a 25-mm round cover slip and then covered with a cellulose acetate dialysis membrane. The membrane had been wetted in distilled water and dried with a filter paper, which exerts a confining pressure due to the hydrophobic effect. An O-ring was placed on the membrane and the entire setup was secured in a Sykes-Moore chamber (Bellco Glass, Cat. #1943-11111). The chamber was then filled with Minimal A medium with 0.2% glucose, 1 mM MgSO_4_, and 20 µg/mL chloramphenicol. Imaging was performed in a temperature-controlled enclosure set to 37 °C. Phase contrast and GFP images were acquired after 1.2 h of growth in the chamber. The microscope and image acquisition were essentially as previously described [60].

### Sucrose density fractionation

For strains carrying MreB^WT^ and MreB^R193C^, inner and outer membranes were separated using a sucrose density gradient as previously described [61, 62]. For DH300 RcsF^WT^ and RcsF^+7^ strains, cell fractionation was adapted from [6]. Four hundred milliliters of cell culture were grown until OD_600_=0.6. Cells were harvested via centrifugation at 6,400*g* and 4 °C for 15 min, washed with TE buffer (50 mM Tris-Cl [pH 7.8], 1 mM EDTA), and resuspended in 20 mL of TE buffer. One milligram of DNase I (Roche), 1 mg of RNase A (Thermo Scientific) and a tablet of Protease Inhibitor Cocktails (Roche cOmplete™) were added to cell suspensions, and cells were passed through a French pressure cell at 12,000 psi. After adding MgCl_2_ to a final concentration of 2 mM, the lysate was centrifuged at 4,200*g* and 4 °C for 8 min to remove cell debris. Then, 16 mL of the supernatant were placed on top of a two-step sucrose gradient (2.3 mL of 2.02 M sucrose in 10 mM HEPES [pH 7.5], 6.6 ml of 0.77 M sucrose in 10 mM HEPES [pH 7.5]). The samples were centrifuged at 130,000*g* for 3 h at 4 °C in a 55.2Ti Beckman rotor. After centrifugation, the soluble and membrane fractions (12 mL) were collected. The membrane fraction was diluted two times with 10 mM HEPES [pH 7.5]. To separate the membranes, 7 mL of the diluted membrane fraction were loaded on top of a second sucrose gradient (10.5 mL of 2.02 M sucrose, 12.5 ml of 1.44 M sucrose, 7 ml of 0.77 M sucrose, all in 10 mM HEPES [pH 7.5]). Samples were then centrifuged at 82,000*g* for 16 h at 10 °C in a SW 28 Beckman rotor. Approximately 30 fractions of 1.5 mL were collected, and odd-numbered fractions were loaded on SDS-PAGE gels, transferred onto a nitrocellulose membrane, and probed with specific antibodies.

### Immunoblotting

To visualize MreB^WT^ and MreB^R193C^ fractionation, protein samples were separated by SDS-PAGE and transferred onto PVDF membranes (IMMOBILON P). To visualize DH300 RcsF^WT^ and DH300 RcsF^+7^ fractionation, protein samples were separated in 4-12% SDS-PAGE gels (Life Technologies) and transferred onto nitrocellulose membranes (GE Healthcare Life Sciences). The membranes were then blocked with 5% skim milk in 50 mM Tris-HCl [pH 7.6], 0.15 M NaCl, and 0.1% Tween20 (TBS-T). TBS-T was used in all subsequent steps of the immunoblotting procedure. Anti-BamA (1:10,000, gift from Lithgow lab at Monash University raised against the soluble BamA POTRA domains [63]), anti-SecG (1:6000, gift from Tokuda lab at University of Morioka, Japan), anti-RcsF (1:20,000 [64]), anti-DsbD (1:2,000 [65]) and anti-Lpp (1:7000) [1] rabbit antisera were used as primary antibodies. The membranes were incubated with horseradish peroxidase-conjugated goat anti-rabbit IgG (Sigma) at a 1:5,000 dilution. Labelled proteins were detected via chemiluminescence (Pierce ECL Western Blotting Substrate, Thermo Scientific) and exposed on X-ray films (Kodak Biomax MR-1) or visualized using GE ImageQuant LAS4000 camera (GE Healthcare Life Sciences).

### Cryo-electron microscopy

MreB^WT^ and MreB^R193C^ strains were grown in LB overnight at 37 °C, diluted to an OD_578_ of 0.0075, and grown in LB at 37 °C until an OD_578_ of 0.2. Cells were collected by centrifugation for 5 min at 1000*g* at 22 °C and concentrated to an OD_578_ of 30 in fresh LB. Concentrated cells (3.5 µL) were immediately applied on glow-discharged (two cycles of 45 s at 15 mA) C-flat^TM^ 4/1 grids (Photochips, Inc). Cells were plunge-frozen using a Vitrobot Mark IV (Thermo Fisher) with a wait time of 0 s, blot time of 3 s, blot force of 3, and drain time of 0 s at constant 100% humidity and at 22 °C. Transmission electron microscopy (TEM) images were collected at the EMBL Electron Microscopy Core Facility using a 200 keV FEI Talos Arctica TEM (Thermo Fisher) equipped with an autoloader and Falcon II direct electron detector (Thermo Fisher) at a pixel size of 0.3266 nm (nominal magnification 45,000X), and a defocus of -10 µm. To image entire cells at high magnification, projection images were collected as a montage of stacks of the field of view in Serial EM software [66].

### Intermembrane distance measurement

Montaged images were pre-blended in etomo [67] and the edges were fixed manually in MIDAS [67]. Membranes were segmented in Fiji [68] after application of a 3-nm Gaussian filter and 10-nm variance filter. Intermembrane distance measurements were carried out using a custom Python script that calculates the distance between two major peaks in the image using a gray-scale gradient. This calculation results in values corresponding to the distance between the centers of the inner and outer membranes, referred to as intermembrane distance. To account for spatial autocorrelation, each intermembrane distance calculation was an average over a 50 nm segment. The poles of the cells were excluded. The outcomes of our analysis were manually validated, and clear miscalculations were removed from the final histograms (Fig. S4B). In total, ∼30 cells from each strain were quantified for each of the three biological replicates.

## Code availability

Scripts for cryo-electron microscopy image analysis are available at the repository https://github.com/martinschorb/membranedist.

## Acknowledgements

The authors thank the Huang and Typas labs for useful discussions. This work was supported by a National Science Foundation Graduate Research Fellowship (to A.M.), an ARCS Fellowship (to A.M.), NIH R01 GM080279 (to M.G.), a James McDonnell Postdoctoral Fellowship (to H.S.), EMBL core funding and a DFG grant (TY 116/2-1) for SPP1617 (to A.T.), NIH Director’s New Innovator Award DP2OD006466 (to K.C.H.), NSF CAREER Award MCB-1149328 and Award EF- 2125383 (to K.C.H.), and the Allen Discovery Center at Stanford on Systems Modeling of Infection (to K.C.H.). K.C.H. is a Chan Zuckerberg Biohub Investigator. This work was also supported in part by the National Science Foundation under Grant PHYS-1066293 and the hospitality of the Aspen Center for Physics.

---

**Figure S1:**
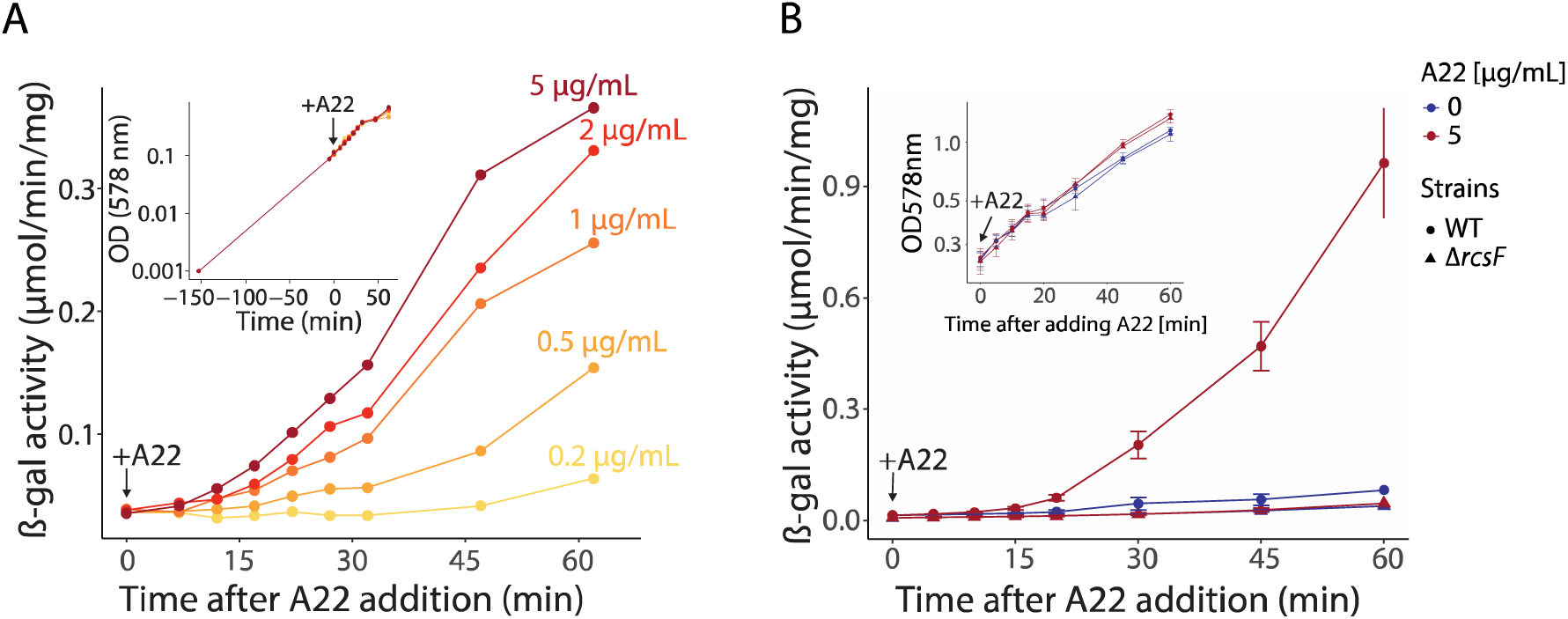
Activation of the Rcs system during A22 treatment is dose-dependent. A) The Rcs system is activated proportionally to the concentration of A22. Activation was measured by monitoring induction of chromosomal *rprA::lacZ*. Cells were treated with A22 at OD_578_=0.3. Inset: growth was unaffected by A22 addition. B) The Rcs system was activated in wild-type but not Δ*rcsF* cells. Data are *n*=4 biological replicates of the experiment in Fig. 1A.

**Figure S2:**
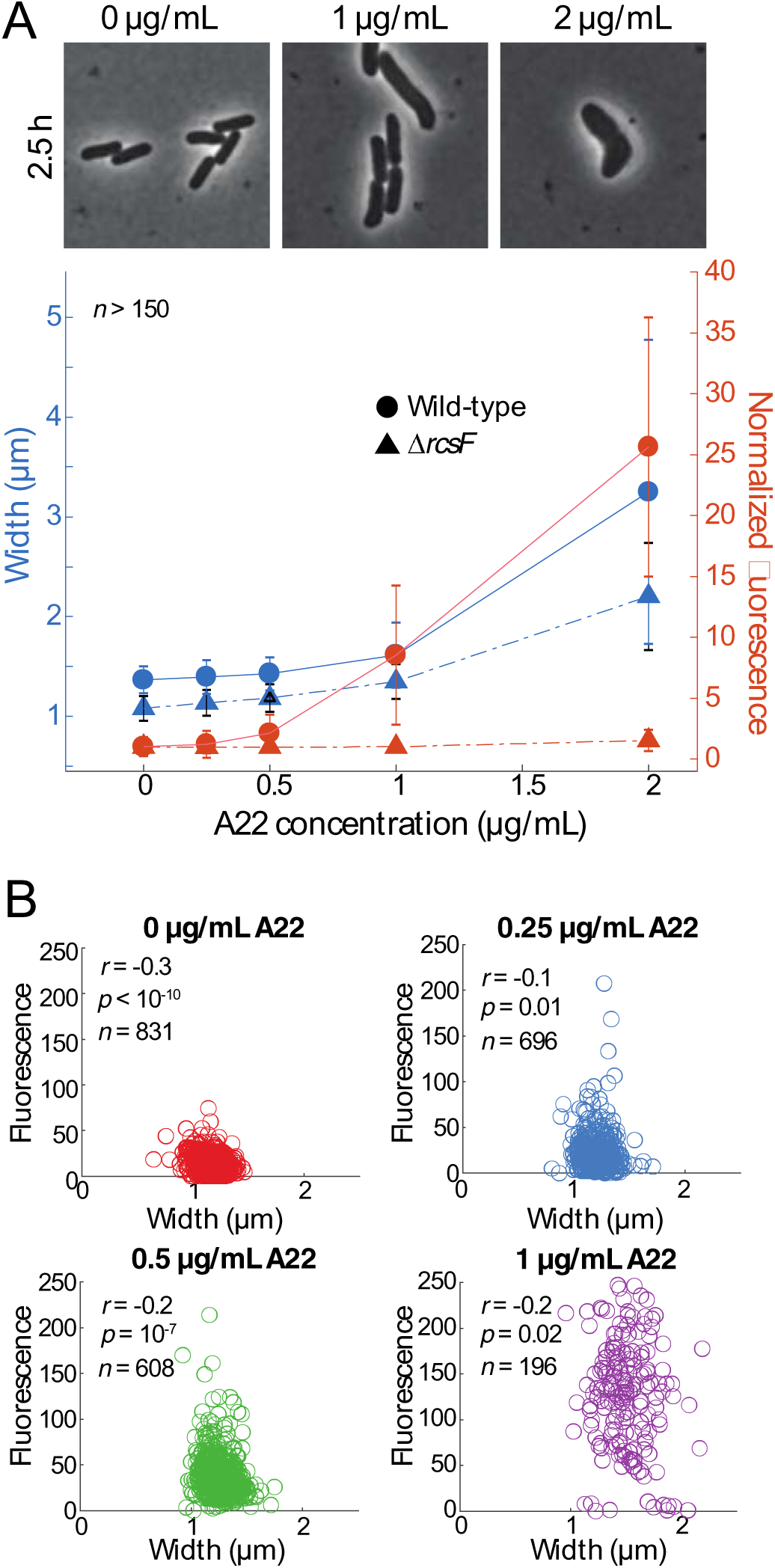
rprA-msfGFP expression is dependent on Rcs activation and is not strongly correlated with natural width variation across a population. A) msfGFP intensity from an *rprA* reporter the pMZ13 as measured via single-cell imaging after 2.5 h of A22 treatment increased with A22 concentration in wild-type but not Δ*rcsF* cells. B) The natural variation in cell width across each population after 2.5 h of A22 treatment was not strongly correlated with msfGFP intensity from the *rprA* promoter on pMZ13.

**Figure S3:**
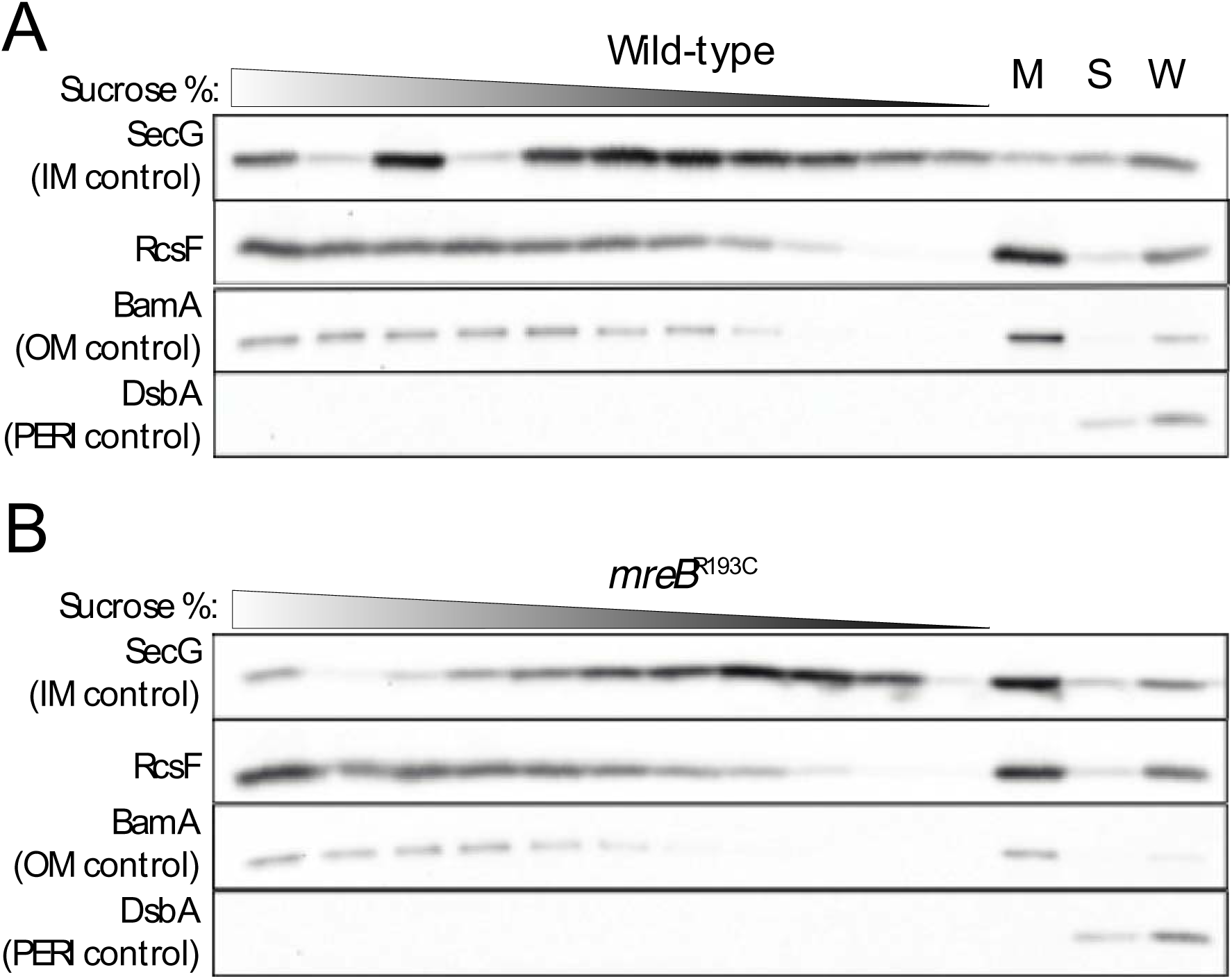
Sucrose-gradient fractionation shows that RcsF is localized in the outer membrane in both wild-type (A) and *mreB^R193C^* (B) cells. Immunoblotting after sucrose-gradient fractionation of RcsF and other control proteins (SecG for the inner membrane, BamA for the outer membrane, and DsbA for the periplasm). OM, outer membrane; IM, inner membrane; PERI, periplasm; M, total membrane sample prior to fractionation; S, soluble non-membrane fraction; W, whole-cell lysate.

**Figure S4:**
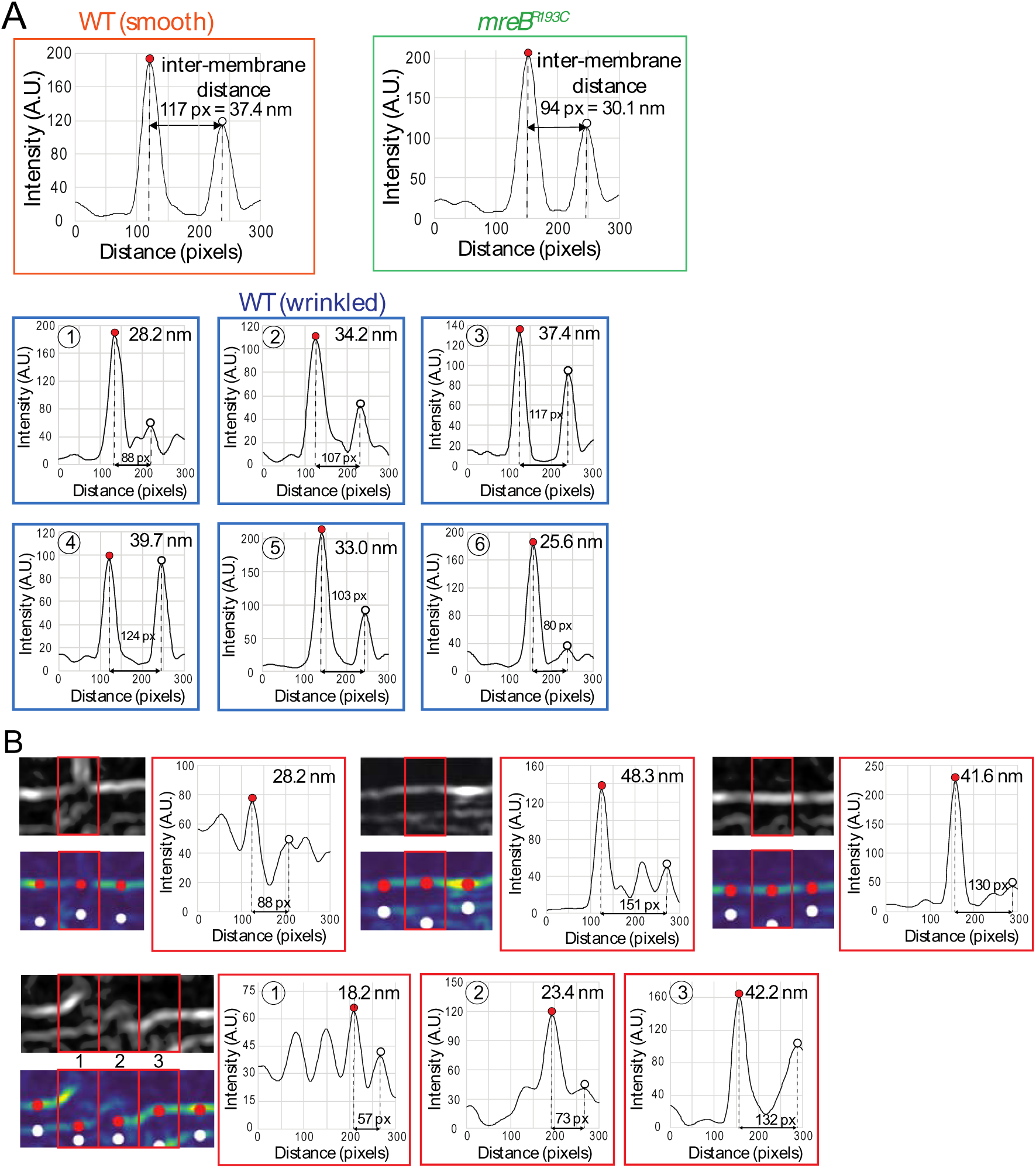
Intermembrane distance is calculated based on the distance between bright cross-sections identified as membranes in processed EM images. A) Peaks were identified in the intensity averaged across a 50-nm segment in the direction perpendicular to the membranes. The two highest peaks correspond to the middle of the outer membrane (left peak, red dot) and inner membrane (right peak, white dot). Shown are examples corresponding to the boxes in Fig. 4D (wild-type straight membrane, orange box; wild-type wrinkled membrane, cyan boxes 1-6; *mreB^R193C^* straight membrane, green box). px, pixel. B) Examples of measurements visually identified as erroneous and hence removed due to multiple reasons. Top, from left to right: membrane was discontinuous, a high-contrast object was located in one membrane, insufficient contrast in one membrane. Bottom: incorrect blending of two cryo-EM fields of view. px, pixel.

**Figure S5:**
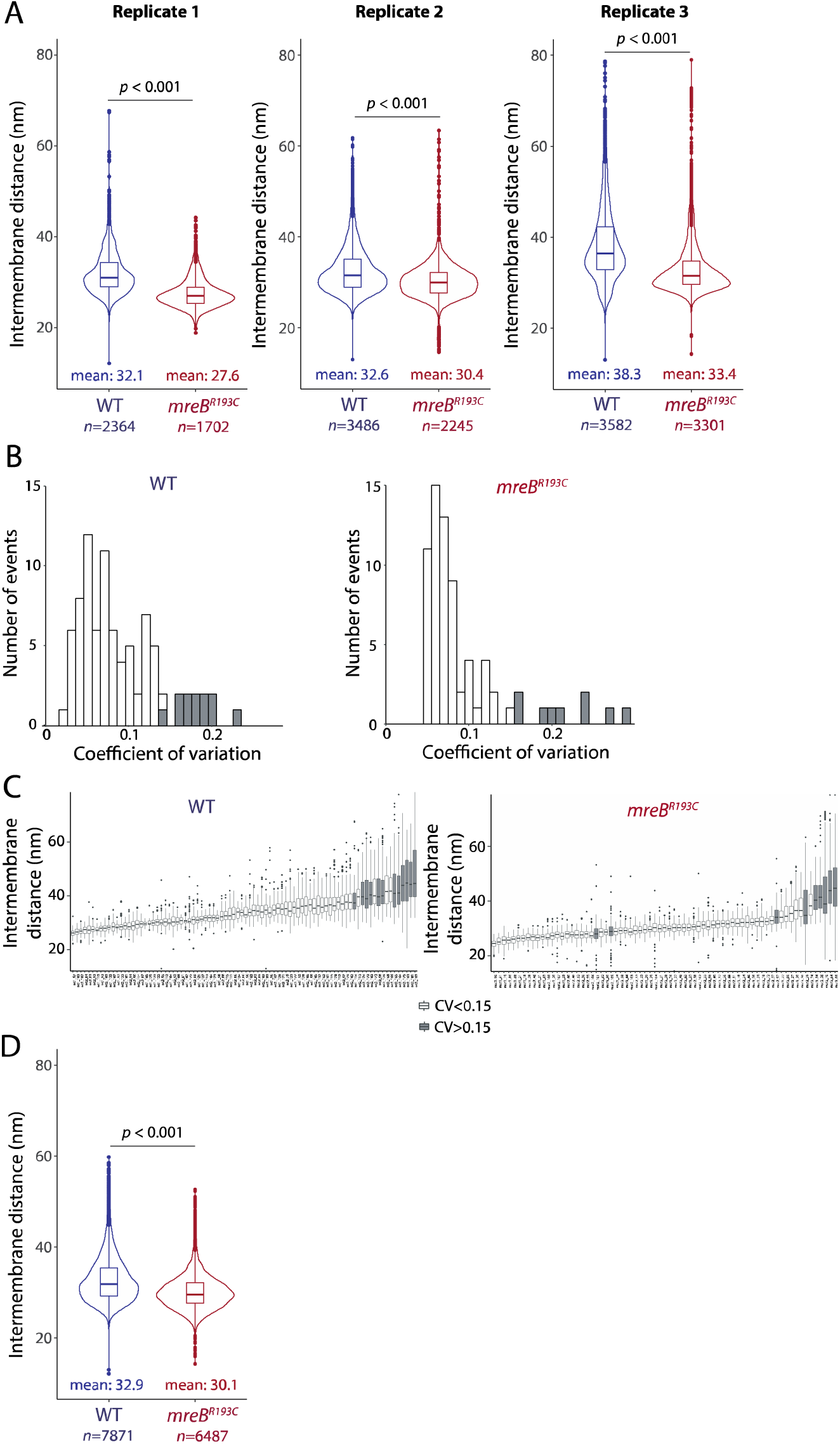
The difference in intermembrane distance between wild-type and *mreB^R193C^* cells is consistent, even though intermembrane distance measurements can vary within a cell, between cells in a population, and between biological replicates. A) Intermembrane distance was consistently higher in wild-type cells compared with *mreB^R193C^* cells across three biological replicates. Means are in nm. *n* represents the number of points at which measurements were made, for ∼30 cells of each strain in each replicate. B) The distribution of coefficients of variation (CV, standard deviation (SD) divided by mean) across cells for wild-type (left) and *mreB^R193C^* (right) cells. Cells with CV>15% are colored in gray. All three biological replicates were included. C) Intermembrane distances within each wild-type (left) and *mreB^R193C^* (right) cell, sorted by mean intermembrane distance. Cells with CV>15% are colored in gray. D) Intermembrane distance was consistently higher in wild-type cells compared with *mreB^R193C^* cells when considering only cells with CV<15%. *n* represents the number of points at which measurements were made, for 74 wild-type and 62 *mreB^R193C^* cells from three biological replicates. Violin plots in (A,D) show the smoothened density of measurements, and boxplots show the median and first and third quartiles.

**Figure S6:**
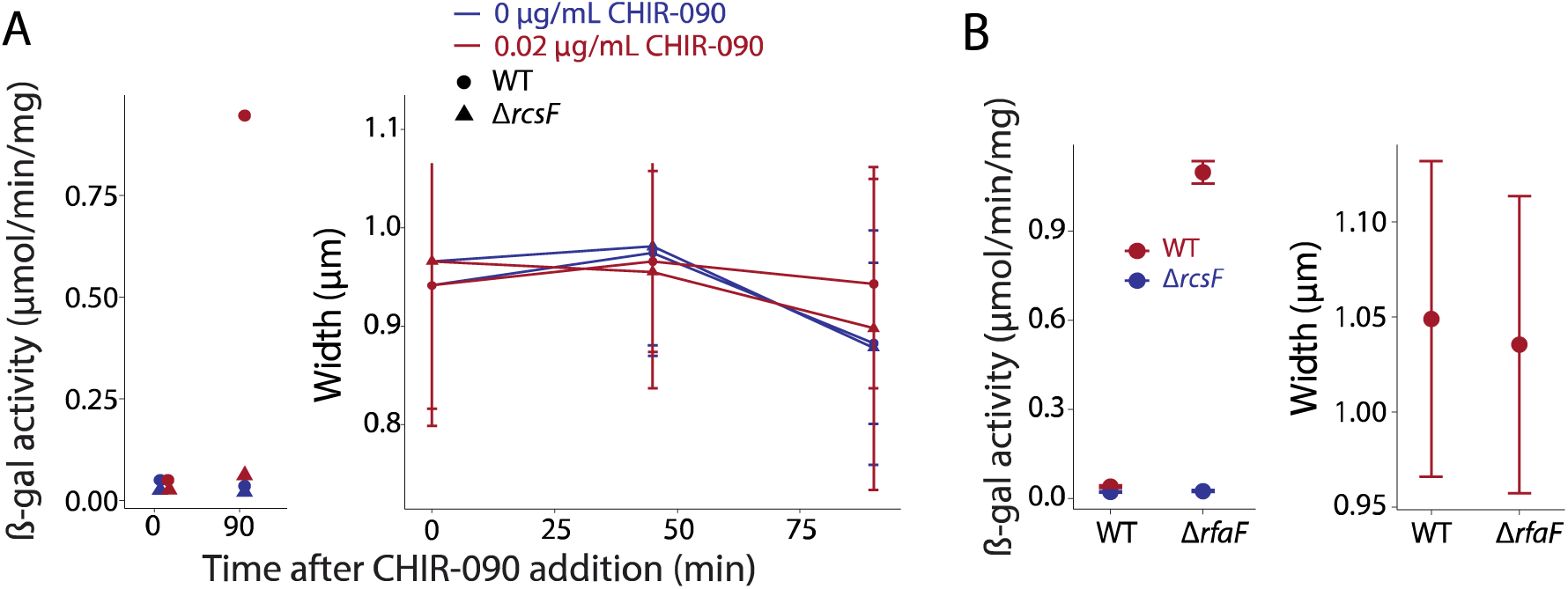
Cell width is not affected by CHIR-090 treatment or deletion of rfaF. A) Treatment with 0.02 µg/mL activates the Rcs pathway (left, *n*=1 replicate) even though cell width is unaffected (right, *n*>200 cells for each data points). B) The Rcs pathway is activated in Δ*rfaF* cells (left, *n*=3 replicates) even though cell width is unaffected (right, *n*=1142 or 926 cells for wild-type (WT) and Δ*rfaF*, respectively). For width measurements in (A,B), data are mean values and error bars represent 1 standard deviation.

**Table S1:** Strains used in this study.

| Strain | Relevant genotype or features | Source |
| --- | --- | --- |
| Aska collection | $\Delta rcsF::cat$ | Mori laboratory (Nara, Japan) |
| DH300 | *rprA::lacZ* MG1655 (*argF-lac*)U169 used as WT | [28] |
| CAG60488 | DH300 $\Delta pgm::kan$ | This study |
| Keio collection of gene deletions | $\Delta rcsF::kan$, $\Delta opgG::kan$, $\Delta opgH::kan$, $\Delta tolB::kan$, $\Delta pgm::kan$ | [14] |
| MG1655 | F- lambda- *ilvG*- *rfb*-50 *rph*-1 | [69] |
| MMR60 | MG1655 $\Delta xylA$ $\Delta xylFG::P_{tetA}::mCherry$ | [56] |
| NT1032 | DH300 $\Delta rcsF::cat$ $\Delta opgG::kan$ | This study |
| NT1033 | DH300 $\Delta rcsF::cat$ $\Delta opgH::kan$ | This study |
| NT1035 | DH300 $\Delta rcsF::cat$ $\Delta tolB::kan$ | This study |
| NT1036 | DH300 $\Delta rcsF::cat$ $\Delta pgm::kan$ | This study |
| NT1038 | DH300 $\Delta opgG::kan$ | This study |
| NT1039 | DH300 $\Delta opgH::kan$ | This study |
| NT1041 | DH300 $\Delta tolB::kan$ | This study |
| NT4043 | DH300 $\Delta rcsF::kan$ | This study |
| RDM893 | MG1655 $\Delta mreBCD$ | [31] |
| NT4151 (MreB<sup>WT</sup>) | RDM893 pMreB<sup>WT</sup> | [31] |
| NT4157 (MreB<sup>R193C</sup>) | RDM893 pMreB<sup>R193C</sup> | [31] |
| KCH1453 | RDM893 $\Delta rcsF$ | This study |

**Table S2:** Plasmids used in this study.

| Plasmid | Features | Source or notes |
| --- | --- | --- |
| pAM238 | IPTG-regulated $P_{lac}$, pSC101-based, spectinomycin | [70] |
| pAC581 |  | [54] |
| pMZ13 | *msfGFP* under *rprA* promoter, pAC581-based, chloramphenicol | This study |
| pSC202 (pRcsF<sup>WT</sup>) | pAM238 with *rscF*<sup>WT</sup> | [6] |
| pRcsF<sup>+7</sup> | pAM238 with *rscF*<sup>+7</sup> | [1] |
| pMreB<sup>WT</sup> |  | [31] |
| pMreB<sup>D78V</sup> |  | [31] |
| pMreB<sup>V316A</sup> |  | [31] |
| pMreB<sup>V236A</sup> |  | [31] |
| pMreB<sup>P314L</sup> |  | [31] |
| pMreB<sup>S10P</sup> |  | [31] |
| pMreB<sup>R193C</sup> |  | [31] |

## Notes

### Competing Interest Statement

The authors have declared no competing interest.

